# Verapamil limits inflammation by restoring VGCC-driven epithelial Ca^2+^ in models of cystic fibrosis

**DOI:** 10.1101/2025.10.12.681885

**Authors:** Sarahdja Cornélie, Bastien Suire, Sylvaine Huc-Brandt, Laure Yatime, Christian M Dupont, Stephen A Renshaw, Georges Lutfalla, R Andres Floto, Céline Cougoule, Audrey Bernut

**Author notes:** correspondence / Laboratory of Pathogen and Host Immunity, CNRS, Inserm, University of Montpellier, 34090 Montpellier, France. **Author Contributions**: AB conceived the study, designed experiments, analyzed data and wrote the manuscript, with input from SC, LY, GL, RAF, SAR, and CC. GL, CC, and AB guided and supervised the work. SC, SH-B, CMD, GL and AB performed zebrafish experiments. BS and CC were responsible for the *in vitro* testing in epithelial cell models. LY performed sequences analyses. All authors contributed to the article and approved the submitted version.

## Abstract

**Rationale:** In cystic fibrosis (CF), caused by mutations in the *CFTR* gene, excessive neutrophilic inflammation drives lung damage and premature mortality. Current anti-inflammatory therapies have limited efficacy, partly due to an incomplete understanding of the mechanisms underlying CF-associated inflammation. Increased epithelial Ca²⁺ signaling has been implicated, but how CFTR dysfunction perturbs the CFTR/Ca²⁺ axis to promote inflammation remains unclear.

**Objectives:** Using Cftr-depleted zebrafish and human approaches, we sought to *i)* define how CFTR dysfunction alters epithelial Ca²⁺ signaling and promotes inflammation, and *ii)* assess whether Ca²⁺ channel antagonists can attenuate inflammatory damage in CF.

**Methods and Results:** We generated a transgenic zebrafish line expressing the Ca²⁺ reporter GCaMP6 under an epithelial promoter to dynamically map epithelial Ca²⁺ activity *in vivo*. Cftr-depleted zebrafish exhibited exaggerated epithelial Ca²⁺ elevations following injury compared with wild-type animals. Genetic or pharmacological inhibition of L-type voltage-gated Ca²⁺ channels (VGCCs) normalized epithelial Ca²⁺ responses and reduced both oxidative stress and neutrophil recruitment. Among FDA-approved VGCCs antagonists tested, only verapamil promoted resolution of neutrophilic inflammation and improved tissue repair in CF zebrafish. Mechanistically, verapamil reduced aberrant epithelial Ca²⁺ fluxes, ROS generation and NF-κB activation. These effects were confirmed in the human CF epithelial cell CFBE41o⁻, where verapamil reduced Ca^2+^ and oxidative stress.

**Conclusions:** These findings establish dysregulated Ca²⁺ signaling *via* L-type VGCCs as a pathogenic driver of inflammation in CF and identify verapamil as a promising therapeutic candidate. By restoring Ca²⁺ and oxidative homeostasis, verapamil alleviates inflammatory damage, supporting its repurposing as a candidate anti-inflammatory therapy in CF.

## Introduction

Cystic fibrosis (CF) is a multisystem genetic disorder caused by biallelic mutations in the CF transmembrane conductance regulator (*CFTR*) gene, that encodes a chloride and bicarbonate channel expressed in epithelial cells (1, 2). In the airways, loss of CFTR function disrupts ion transport, leading to dehydrated mucus and compromised mucociliary clearance, creating an environment predisposing to recurrent and chronic infections. These conditions drive an excessive and persistent neutrophil-dominated response that starts early in life, fails to eradicate pathogens, and causes progressive airway injury culminating in fatal respiratory failure.

Inflammation in CF, once considered a secondary consequence of infections, is now recognized as being amplified by intrinsic defects in epithelial and immune cells linked to CFTR dysfunction. In CF, airway epithelia exhibit elevated basal expression of pro-inflammatory cytokines and a hyperresponsive phenotype upon injury or microbial stimulation, driven in part by oxidative stress and aberrant activation of NF-κB and inflammasome, which collectively promote an excessive neutrophil response (3). In parallel, CFTR-deficient neutrophils fail to undergo apoptosis and reverse migration (4, 5), enhancing and maintaining their pro-inflammatory activity within the airways.

Conventional anti-inflammatory therapies such as corticosteroids or high-dose ibuprofen have limited efficacy and significant side effects (6). CFTR modulators, including the triple-combination elexacaftor–tezacaftor–ivacaftor, provide partial anti-inflammatory benefit but fail to fully normalize airway inflammation, particularly in patients with advanced disease or those ineligible for modulators (7). These limitations highlight the urgent need for new therapeutic strategies targeting inflammatory pathology in people with CF.

Emerging evidence implicates dysregulated epithelial calcium (Ca²⁺) signaling in the initiation and persistence of CF-related inflammation. In healthy airway epithelia, intracellular Ca²⁺ regulates mucin secretion, ciliary beating, as well as inflammatory and repair processes (8). However, in CF, loss of CFTR disrupts epithelial Ca²⁺ homeostasis *via* increased store-operated Ca²⁺ entry (SOCE), enhanced transient receptor potential (TRP) activity, and impaired mitochondrial Ca²⁺ extrusion, leading to sustained intracellular Ca²⁺ elevation (9). This aberrant signaling promotes an increased generation of reactive oxygen species (ROS) and exaggerated pro-inflammatory cytokines release. Yet, most evidence derives from *in vitro* or *ex vivo* studies, and *in vivo* mechanistic data directly linking CFTR dysfunction, Ca²⁺ dysregulation, and inflammation, especially neutrophil response, are lacking.

Here, using Cftr-depleted zebrafish, a clinically relevant vertebrate model recapitulating key aspects of CF immune-pathogenesis, along with human bronchial epithelial cells, we confirm a mechanistic link between epithelial Ca²⁺ dysregulation and CF-related neutrophilic inflammation. In particular, we *i)* identify L-type voltage-gated Ca²⁺ channels (VGCCs) as critical contributor to this process and *ii)* demonstrate that verapamil, an FDA-approved L-type VGCC blocker, by normalizing epithelial Ca²⁺ fluxes, reduces oxidative stress and NF-kB activity, attenuates neutrophil recruitment and improves tissue repair. These findings position L-type VGCC inhibition as a promising therapeutic strategy to limit airway inflammation and tissue damage in CF.

## Methods

Human cells, zebrafish lines, and detailed methods associated with the procedures described in this study are available in the data supplement.

## Results

### Loss of CFTR leads to an abnormal elevation of epithelial Ca^2+^ fluxes in injured tissues *in vivo*

To interrogate the relationship between CFTR function, epithelial Ca^2+^ signaling and inflammatory responses, we first compared the characteristic pattern of epithelial Ca^2+^ flux, induced by inflammatory stimuli, in normal *vs* Cftr-depleted zebrafish (using *cftr* morphants (c*ftr* MO) (10)). To do so, we generated a transgenic line expressing the genetically encoded Ca²⁺ indicator GCaMP6 under the control of the epithelial *krtt1c19* promoter (**Figure 1A**), enabling direct visualization of epithelial Ca²⁺ dynamics. Mechanical tissue injury (4), which triggers rapid rise in cytoplasmic Ca²⁺ level in epithelial cells (11, 12), was used to initiate an inflammatory response. In Cftr-depleted zebrafish, fluorescence analyses revealed aberrant Ca²⁺ activity characterized by an exaggerated early elevation and sustained Ca²⁺ levels at the injury site (**Figures 1B-C; S1A-B**). Comparable results were obtained using the dye-based Ca²⁺ reporter Fluo3-AM (**Figures S1C-D**). These results are analogous to the elevated intracellular Ca²⁺ levels reported in CF epithelial cells compared to non-CF cells (13–15).

**Figure 1:**
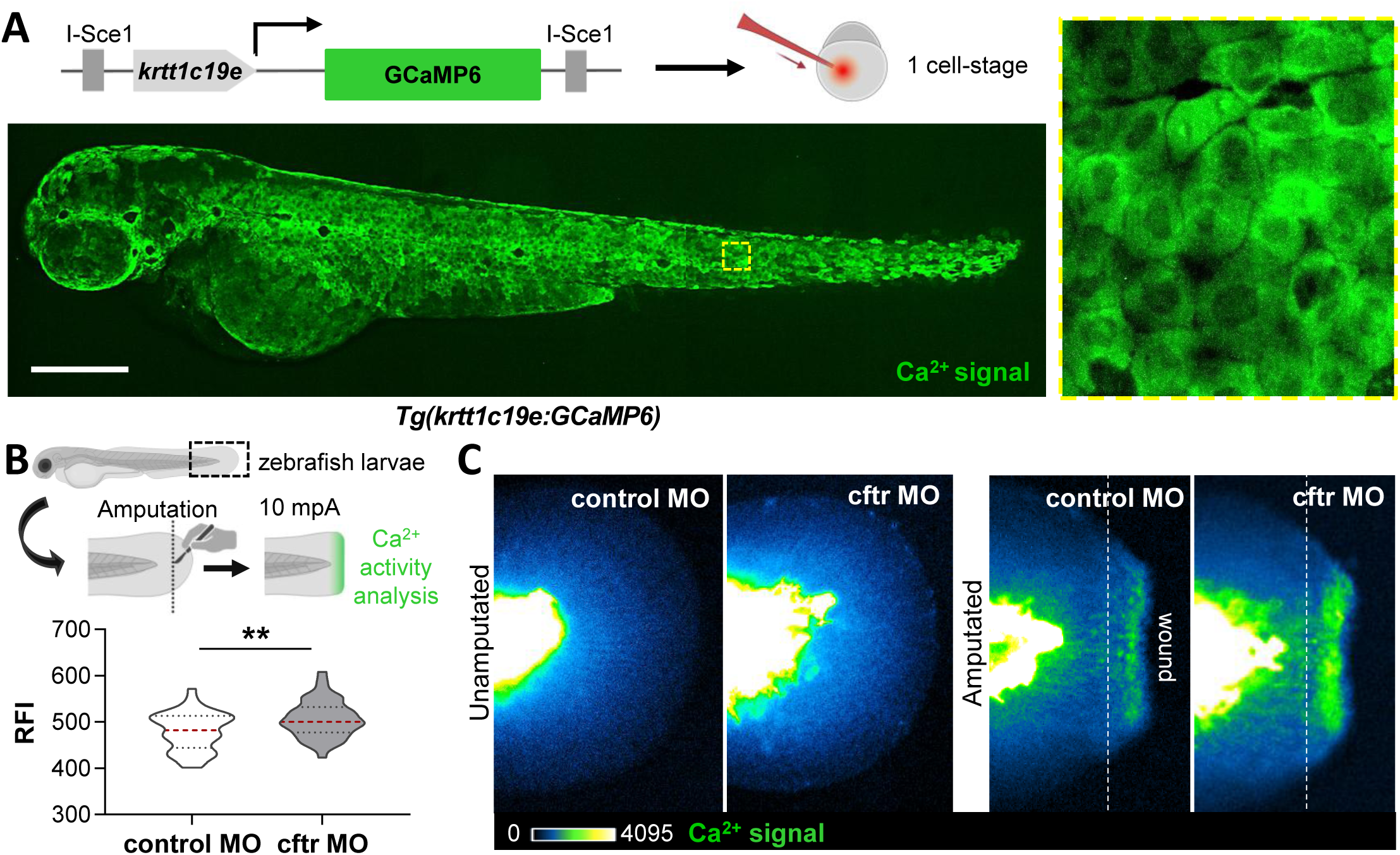
Injury-induced epithelial Ca^2+^ fluxes are exacerbated in Cftr-depleted zebrafish. **(A)** Generation of a transgenic line expressing the Ca^2+^ indicator GCaMP6 in epithelial cells (*Tg(krtt1c19e:GCaMP6)ump14Tg*) using the I-Sce1 meganuclease system. In this transgenic zebrafish, epithelial promoter krttc19e drives the expression of GCaMP6 in epithelial cells. Representative photomicrographs of the overall pattern of fluorescence of a *Tg(krtt1c19e:GCaMP6)ump14Tg* larvae exposed to ionomycin (3 days post-fertilization (dpf); scale bar: 200 µm). **(B-C)** Control (control MO) and *cftr* morphants (*cftr* MO) *Tg(krtt1c19e:GCaMP6))ump14Tg* were tail fin amputated, then epithelial Ca^2+^ activity at wound sites was measured and analyzed at 10 min post-amputation (mpA). Quantitative fluorescence microscopy analysis of epithelial Ca^+2^ fluxes quantified by relative fluorescence intensity (RFI) (n=45 from three independent experiments; mean and quartiles shown, Student T-test) **(B)** and associated pseudocolored photomicrographs of uninjured and injured tails revealing epithelial Ca^2+^ activity **(C)**.

These findings confirm that loss of CFTR function disrupts cytoplasmic Ca^2+^ levels in epithelial cells, supporting the idea that CFTR plays an important role in the regulation of epithelial Ca^2+^ signaling.

### L-type Ca^2+^ channels drive neutrophilic inflammation in Cftr-deficient zebrafish

Since CFTR dysfunction causes abnormal Ca^2+^ levels in epithelial cells, we next sought to *i)* identify the Ca^2+^ channels involved in increased Ca^2+^ fluxes to injury in CF animals, and *ii)* determine whether Ca^2+^ signaling regulates inflammatory responses, and how this altered axis might be involved in CF-related inflammation. To do so, we used pharmacological antagonists targeting different channels expressed in epithelia: capsazepin to act against TRP, Cpd5J-4 to inhibit SOCE and nifedipine to block L-type voltage-gated Ca²⁺ channels (VGCC) (**Figure 2A**), assessing their effects on epithelial Ca²⁺ responses to injury. All antagonists significantly reduced injury-induced Ca²⁺ flux in Cftr-depleted larvae (**Figures 2B-C**). While the roles of SOCE and TRP channels in disrupted Ca²⁺ signaling in CF are already well established (9), these results highlight L-type VGCCs as additional, previously unrecognized contributors to aberrant epithelial Ca²⁺ activity in this disease.

**Figure 2:**
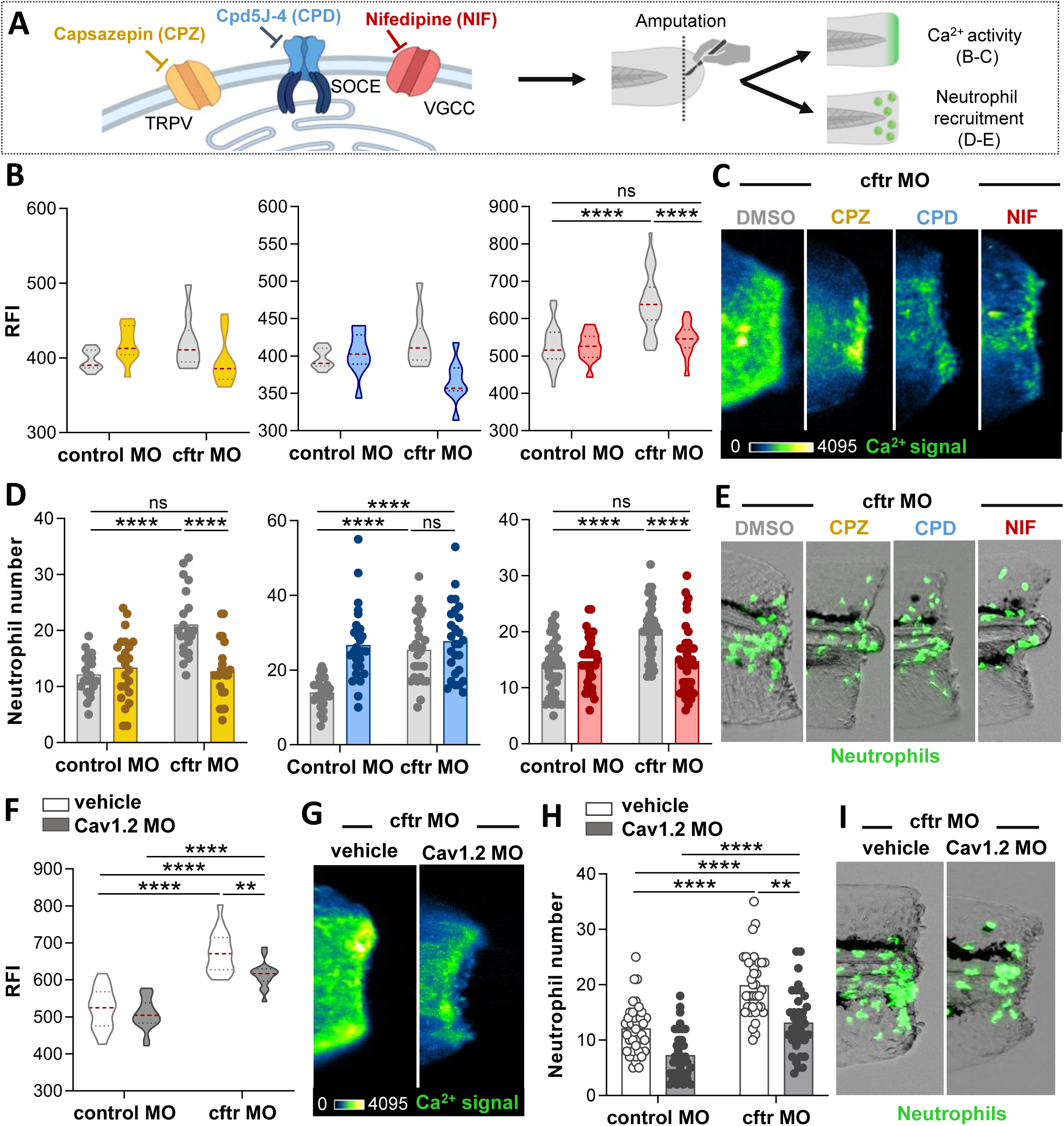
Blockade of L-type Cav1.2 channels reduces Ca²⁺-mediated neutrophilic inflammation in Cftr-deficient zebrafish. **(A-E)** *Tg(krtt1c19e:GCaMP6)ump14Tg* (B-C) or *TgBAC(mpx:eGFP)i114* (D-E) injected with control or *cftr* morpholinos were pretreated with capsazepin (CPZ; yellow), Cpd5J-4 (CPD, blue), nifedipine (NIF, red) or DMSO (vehicle control, grey) prior caudal fin amputation. Larvae are then injured and immediately put back in treatments until analyses. **(B-C)** Epithelial Ca^2+^ activity was observed and analyzed at the wound sites at 20 mpA. Quantification of relative epithelial Ca^2+^ intensity (n=27-31 from 3 independent experiments; Two-way ANOVA, Tukey’s multiple comparisons test) **(B)** and representative pseudocolored photomicrographs of wound-associated Ca^2+^ fluxes in CF zebrafish **(C)**. **(D-E)** Neutrophilic inflammation observed and analyzed at 3 hours post amputation (hpA). Number of neutrophils mobilized (n=35-38 from 3 independent experiments; Two-way ANOVA, Tukey’s multiple comparisons test) **(D)** and representative photomicrographs showing neutrophil accumulation at the wound site in CF zebrafish **(E)**. **(F-G)** Control MO, *cftr* MO, *Cav1.2* MO and double *cftr* MO/ *Cav1.2* MO *Tg(krtt1c19e:GCaMP6)ump14Tg* were caudal fin amputated, and the epithelial Ca^2+^ activity was observed and analyzed at 10 mpA. Relative epithelial Ca^2+^ intensity (n=16-20 from 2 independent experiments; Two-way ANOVA, Tukey’s multiple comparisons test) **(F)** and associated pseudocolored photomicrographs of injured tails revealing epithelial Ca^2+^ activity in *cftr* MO and double *cftr* MO/ *Cav1.2* MO) **(G)**. **(H-I)** Control MO, *cftr* MO, *Cav1.2* MO and double *cftr/Cav1.2* MO *TgBAC(mpx:eGFP)i114* were caudal fin amputated, then the number of neutrophils mobilized at the wound was observed and counted at 3hpA. Number of neutrophils mobilized (n=38 from 3 independent experiments; Two-way ANOVA, Tukey’s multiple comparisons test) **(H)** and representative photomicrographs showing neutrophil accumulation at the wound sites in cftr MO and double *cftr/Cav1.2* MO **(I)**.

We previously showed that loss of Cftr leads to an exuberant neutrophil response to tissue injury in zebrafish (4). Using the *TgBAC(mpx:EGFP)i114* reporter line harboring green fluorescent neutrophils (16), we examined whether Ca^2+^ channels blockade influences CF-related neutrophilic inflammation. As indicated in **Figures 2D-E**, pharmacological inhibition of epithelial Ca^2+^ fluxes from L-type VGCCs markedly reduced neutrophil trafficking to wounds in CF zebrafish.

Importantly, since nifedipine is known to affect neutrophil functions by a non-Ca^2+^-mediated mechanism (17), we sought to confirm that its effect on CF-related neutrophilic inflammation was due to epithelial Ca²⁺ modulation rather than compromised neutrophil chemotaxis. To do so, we used the well-established *in vivo* fMLP microinjection assay to study neutrophil chemotaxis capacity in zebrafish larvae (4). As shown in **Figure S2**, fMLP-stimulated neutrophil chemotaxis assay revealed that nifedipine-treated larvae exhibited a neutrophil recruitment indistinguishable from that of their control counterparts, confirming that nifedipine does not impair neutrophil chemotaxis *per se* in zebrafish larvae.

Having identified L-type VGCCs as an unrecognized pathway of Ca²⁺ dysregulation in CF, we next investigated which specific channels drive the elevated epithelial Ca²⁺ activity in CF zebrafish. Four isoforms of L-type VGCCs have been described in human, Cav1.1, Cav1.2, Cav1.3 and Cav1.4. Also identified in zebrafish (Cav1.1a and 1.1b, Cav1.2, Cav1.3a and 1.3b, Cav1.4a and 1.4b), the human and zebrafish Cav1s are highly conserved (80% homology) (**Figure S3**). Since Cav1.2 is the major L-type VGCC expressed at epithelial surfaces, including lung tissue (18), we were interested in whether this Ca^2+^ channel influenced epithelial Ca^2+^ response during inflammation in zebrafish. While qRT-PCR analysis showed similar expression levels of *Cav1.2* in normal and Cftr-depleted zebrafish (**Figure S4A**), our analysis revealed that genetic knocked-down of *Cav1.2* in CF animals significatively reduced both epithelial Ca^2+^ flux (**Figures 2F-G**) and neutrophil movement (**Figures 2H-I and S4B**) towards injury, suggesting a role for epithelial Ca^2+^ signaling through Cav1.2 in CF-related neutrophilic inflammation.

Together, these findings establish that CFTR deficiency promotes a dysregulated epithelial Ca²⁺ signaling that contributes to neutrophilic inflammation and reveal an underrecognized link between altered L-type VGCC-mediated Ca²⁺ signaling and inflammation in CF.

### Verapamil reduces Ca^2+^-mediated inflammation and tissue damage in Cftr-depleted zebrafish

Excessive, non-resolving neutrophilic inflammation is a hallmark of CF pathogenesis, driving tissue damage and premature mortality. Our data indicate that overactive neutrophil responses in CF larvae arises from abnormal increases epithelial Ca^2+^ elevation, notably *via* L-type VGCC like Cav1.2, since inhibition of these channels can block this downstream effect. This suggests that L-type VGCCs might represent promising therapeutic targets to reduce inflammation in CF.

To assess their therapeutic potential, we evaluated three FDA-approved L-type VGCC antagonists, verapamil, diltiazem, and amlodipine in CF zebrafish (**Figure 3A**). While all three reduced overactive neutrophil mobilization (**Figures 3B and S5**), only verapamil was able to accelerate resolution of inflammation in CF animals, as neither diltiazem nor amlodipine decreased the persistence of wound-associated neutrophils (**Figures 3C-D**). Moreover, although Cftr-depleted zebrafish exhibit impaired tissue repair and abnormal remodeling after tail fin amputation (4, 19), we found that only verapamil-treated animals displayed enhanced tissue repair and reduced tissue damage (**Figures 3E-F**).

**Figure 3:**
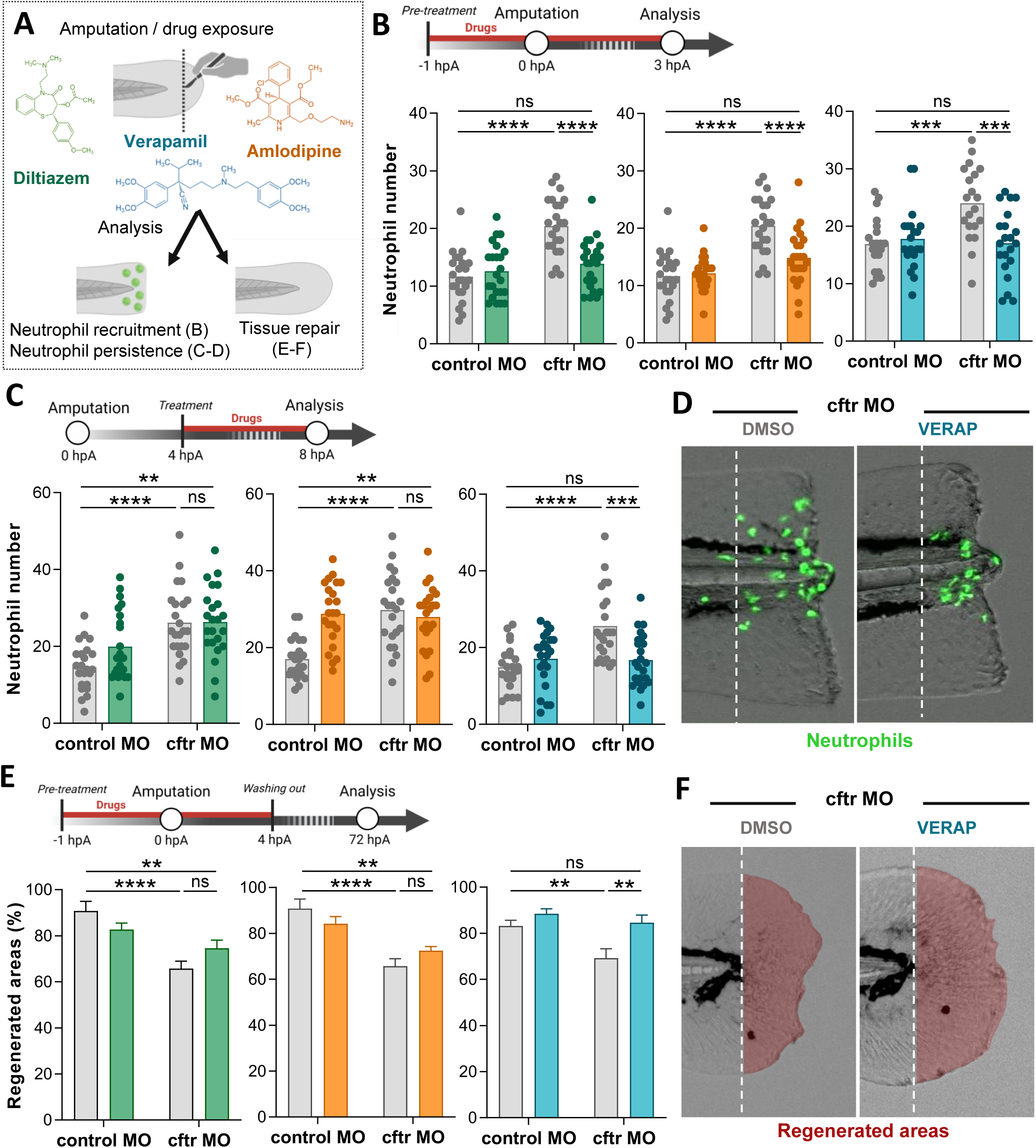
Verapamil exposure alleviates neutrophilic inflammation and tissues damage *in vivo*. **(A)** The effects of diltiazem (green), amlodipine (orange) and verapamil (blue) on neutrophilic inflammation **(B-D)** and tissue repair **(E-F)** were evaluated in zebrafish larvae. **(B)** *TgBAC(mpx:eGFP)i114* injected with control or *cftr* morpholinos were pretreated with drugs or DMSO prior caudal fin amputation. Number of neutrophils mobilized at the wounds at 3 hpA (n=20-23, from 3 independent experiments; Two-way ANOVA, Tukey’s multiple comparisons test). **(C-D)** Control MO and *cftr* MO *TgBAC(mpx:eGFP)i114* were tail fin amputated and treated with drugs or DMSO from 4 hpA. Number of neutrophils at the wounds at 8 hpA (n=23 from 3 independent experiments; Two-way ANOVA, Tukey’s multiple comparisons test) **(C)** and representative photomicrographs of wound-associated neutrophils in CF larvae treated with DMSO or verapamil (VERAP) **(D)**. **(E-F)** Control MO and *cftr* MO were pretreated with drug or DMSO prior caudal fin amputation procedure, then injured and immediately put back in treatments for 4h. The potential of tissue repair was evaluated at 72 hpA. Measurement of regenerated caudal fin areas (n=20 from 3 independent experiments; Two-way ANOVA, Tukey’s multiple comparisons test) **(E)** and brightfield photomicrographs of injured tails revealing fin regrowth in *cftr* MO treated with DMSO or verapamil **(F).**

Pulmonary infections are major features in CF, where they exacerbate neutrophil-dominated inflammation and accelerate progressive tissue damage in the lung. Given its potential, we next evaluated verapamil efficacy during infections with CF-relevant pathogens (*Staphylococcus aureus, Pseudomonas aeruginosa* and *Mycobacterium abscessus*). As expected, bacterial challenge exacerbated neutrophil recruitment, tissue damage and defective repair phenotype in Cftr-depleted zebrafish (**Figure S6A-D**). Importantly, while *in vitro* assays confirmed that verapamil did not affect bacterial growth (**Table 1**), we found that verapamil reduced hyper-neutrophilia and improved tissue repair in infected CF larvae (**Figures S7A-B**).

We next decided to better understand the mechanisms by which verapamil reduces neutrophilic inflammation in CF animals (**Figure 4A**). First, we confirmed that verapamil suppressed injury-induced epithelial Ca²⁺ hyperactivity in Cftr-depleted zebrafish (**Figures 4B-C**). Epithelial Ca²⁺ elevations drive H₂O₂ production, a potent neutrophil chemoattractant (11). Since exaggerated production of ROS by epithelia contributes to neutrophilic inflammation in CF (4), we next proceeded to examine whether verapamil could mitigate oxidative stress in CF larvae. As shown in **Figures 4D-E**, verapamil significantly reduced wound-associated ROS in CF fish compared to DMSO-treated animals. Excessive nuclear factor-κB (NF-κB) pathway activation, another key driver of neutrophil chemotaxis, has been described in the pathology of lung inflammation in CF (20), and was confirmed in zebrafish, with higher Nf-κB signal at wound in Cftr-depleted zebrafish compared to control animals (**Figures S8A-C**). Similar to L-type channel inhibition, verapamil can directly inhibit the NF-κB signaling pathway (21). Therefore, we explored the potential of verapamil to inhibit NF-κB transcription, using a promoter transgenic line. As anticipated, we found that Nf-κB transcription was markedly reduced following verapamil exposure (**Figures 4F-G**). By contrast, while some reports have highlighted the capacity of verapamil to reduce IL1b release, our results showed that verapamil exposure was not sufficient to reduced Il1b transcription in *cftr* MO (**Figures 4H-I**), suggesting that its anti-inflammatory effect does not rely on IL1b inhibition.

**Figure 4:**
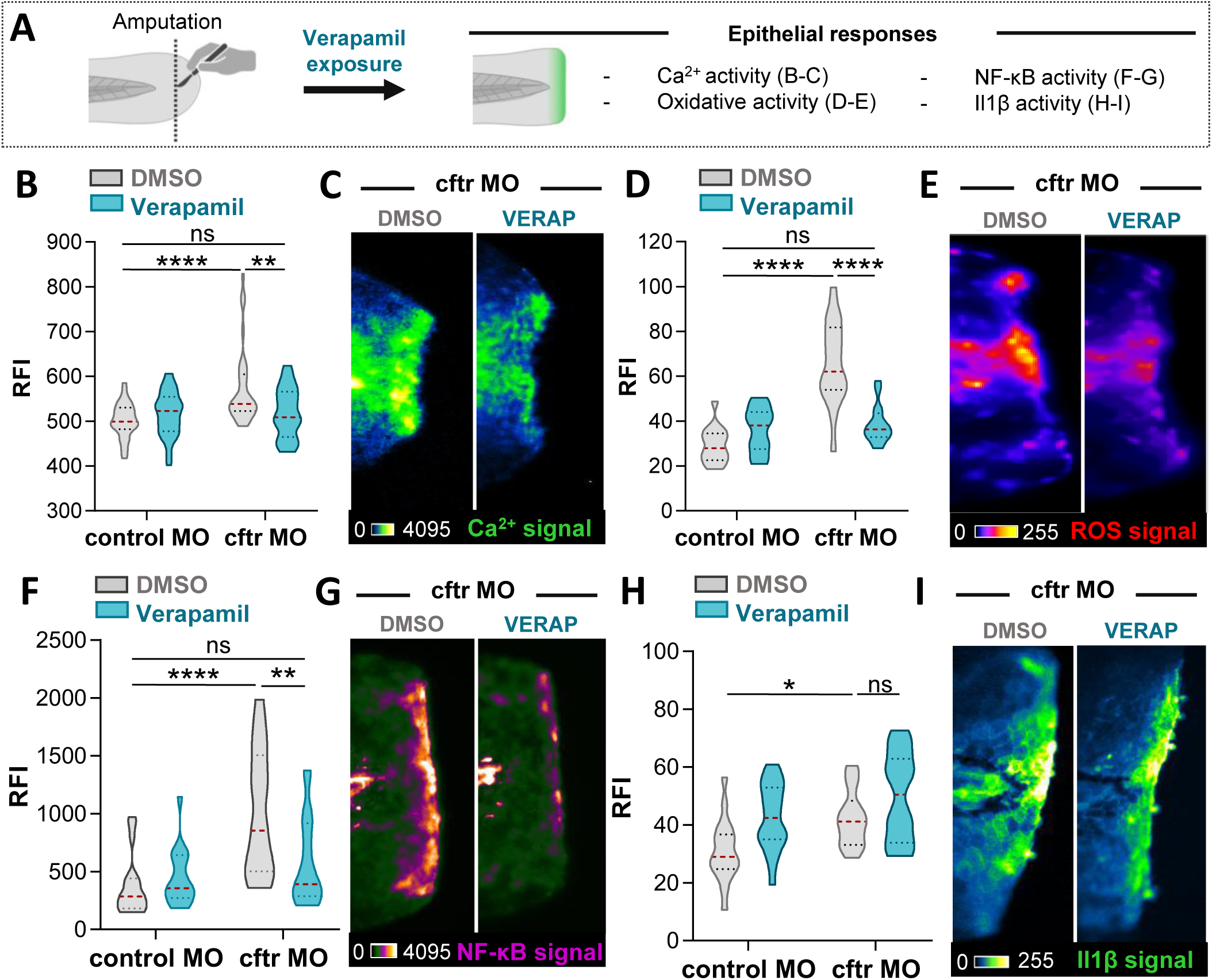
Verapamil reduces epithelial oxidative stress and NF-kB activity in Cftr-depleted zebrafish. **(A)** The effects of verapamil on epithelial responses **(B-I)** were evaluated in zebrafish larvae. **(B-C)** *Tg(krtt1c19e:GCaMP6)ump14Tg* injected with control or *cftr* morpholinos were pretreated with verapamil or DMSO prior caudal fin amputation. Larvae are then injured and immediately put back in treatments until analyses. **(B-C)** Epithelial Ca^2+^ activity was observed and analyzed at the wound sites at 20 mpA. Quantification of relative epithelial Ca^2+^ intensity (n=30 from 3 independent experiments; Two-way ANOVA, Tukey’s multiple comparisons test) **(B)** and representative pseudocolored photomicrographs of wound-associated Ca^2+^ fluxes in CF zebrafish **(C)**. **(D-E)** To label and analyze ROS production, zebrafish were stained with CellROX. Control MO and *cftr* MO were pretreated with verapamil or DMSO before tail fin amputation procedure, then injured and immediately put back in treatments. Relative ROS intensity at 30 mpA (n=20 from 3 independent experiments; Two-way ANOVA, Tukey’s multiple comparisons test) **(D)** and associated pseudocolored photomicrographs of injured tails revealing ROS production in *cftr* MO treated with DMSO or verapamil **(E)**. **(F-G)** Control MO and *cftr* MO *Tg(pNF-kB:EGFP)sh235* were pretreated with verapamil or DMSO prior caudal fin amputation procedure, then injured and immediately put back in treatments. Relative NF-kB intensity at 3 hpA in control MO and *cftr* MO treated or not with verapamil (n=18 from 3 independent experiments; Two-way ANOVA, Tukey’s multiple comparisons test) **(F)** and associated pseudocolored photomicrographs of injured tails revealing wound-associated Nf-κB activity in *cftr* MO treated with DMSO or verapamil **(G)**. **(H-I)** Control MO and *cftr* MO *Tg(il1b:eGFPF)ump3tg* were pretreated with verapamil or DMSO prior caudal fin amputation procedure, then injured and immediately put back in treatments. Relative Il1b intensity at 2 hpA in control MO and *cftr* MO treated or not with verapamil (n=18 from 3 independent experiments; Two-way ANOVA, Tukey’s multiple comparisons test) **(H)** and associated pseudocolored photomicrographs of injured tails revealing wound-associated Il1b activity in *cftr* MO treated with DMSO or verapamil **(I)**.

Together, these findings identify verapamil as a potent modulator of epithelial Ca²⁺ signaling in CF. By restoring Ca²⁺ homeostasis, verapamil limits ROS overproduction and excessive Nf-κB activation, thereby alleviating neutrophilic inflammation and tissue damage in Cftr-deficient zebrafish.

### Verapamil reduces oxidative stress in human CF bronchial epithelial cells

Finally, to validate the efficacy of verapamil in reducing CF-related inflammation in a human system, we examined its effect in the F508del-CFTR human CF bronchial epithelial cell line (CFBE41o-, CFBE), derived from a CF patient (**Figure 5A**). Compared with normal bronchial epithelial cells (16HBE14o-, HBE), CFBE cells displayed exaggerated Ca²⁺ flux, as assessed by Fluo-3 staining (**Figure 5B**). Verapamil treatment efficiently restored Ca^2+^ activity in CFBE to levels comparable with HBE cells. Because oxidative stress is a central driver of neutrophilic inflammation in CF airways, we next assessed the impact of verapamil on ROS production using CellROX staining. Comparative analysis revealed that verapamil-treated CFBE cells exhibited significantly reduced ROS levels relative to DMSO controls (**Figure 5C**).

**Figure 5:**
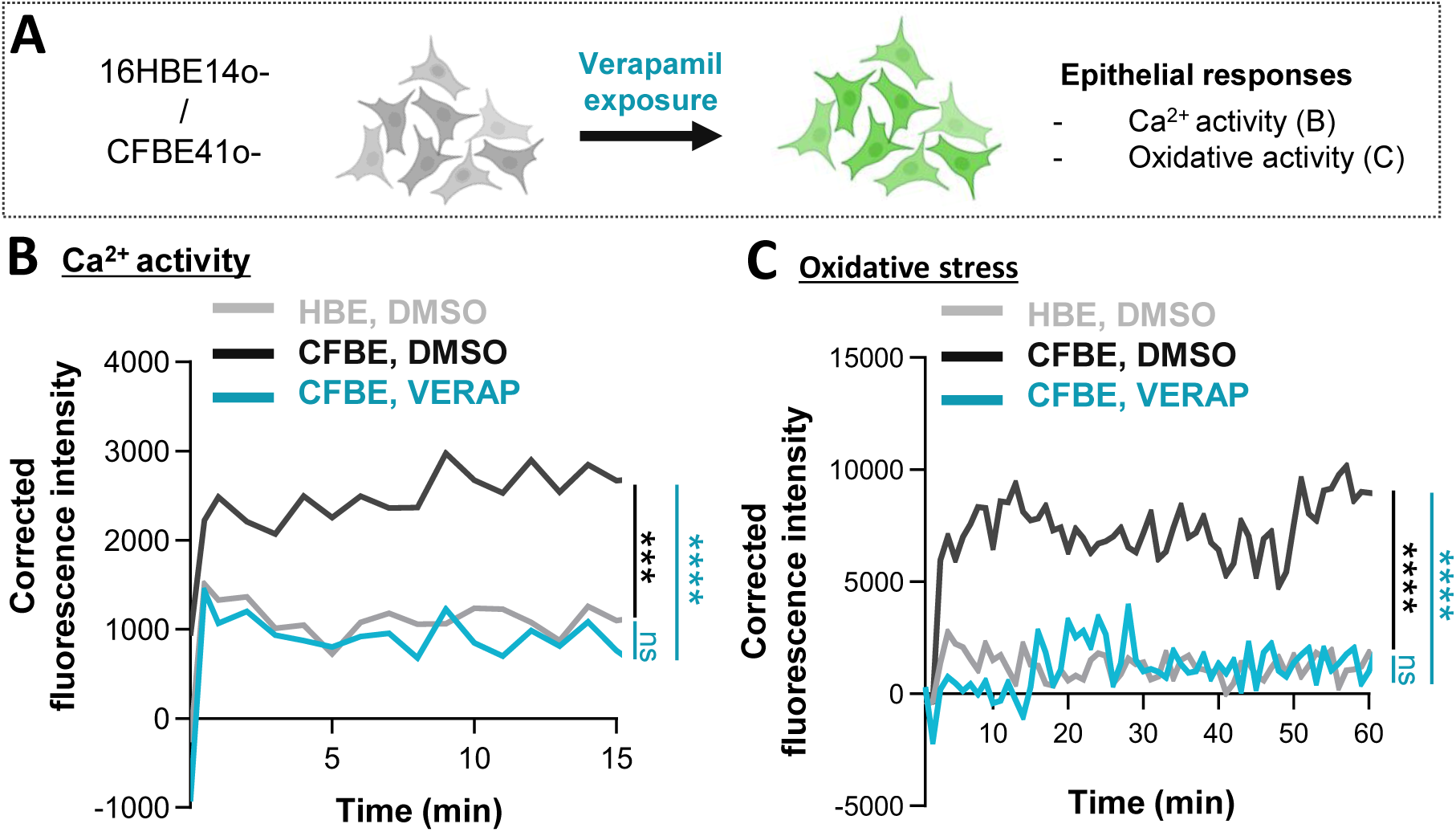
Verapamil reduces excessive Ca^2+^ activity and oxidative stress in human CF bronchial epithelial cells. **(A)** The effects of verapamil on Ca^2+^ fluxes **(B)** and oxidative responses **(C)** were evaluated in CFBE41o- and 16HBE14o-cell lines. **(B)** Epithelial cells were stained with Fluo3-AM to label and analyze Ca^2+^ activity. Epithelial cells were treated with DMSO or verapamil, stimulated with ionomycin, then Ca^2+^ intensity was quantified over 15 min (linear regression, from 4 replicates). **(C)** Epithelial cells were stained with CellROX to label and analyze ROS production. Epithelial cells were treated with DMSO or verapamil, stimulated with cumene hydroperoxide, then ROS intensity was quantified over 60 min (from 4 independent replicates, Two-way ANOVA with Tukey’s multiple comparisons test).

These findings extend our *in vivo* observations and demonstrate that verapamil normalizes epithelial Ca²⁺ signaling and reduces oxidative stress in human CF bronchial epithelial cells, supporting its potential as a therapeutic strategy to limit pulmonary inflammation in CF.

## Discussion

Excessive neutrophil-dominated inflammation is a major contributor of bronchiectasis, airway damage, and ultimately, fatal respiratory failure in individuals with CF. Although CFTR modulators, including the triple-combination elexacaftor–tezacaftor–ivacaftor, have markedly improved outcomes for many patients, persistent airway inflammation, especially in those with advanced disease or ineligible for modulators, remains an unmet therapeutic challenge.

Among the pathways implicated in CF-related inflammation, dysregulated epithelial Ca²⁺ signaling has emerged as a promising target (9). Pharmacological modulation of Ca²⁺ flux reduces pro-inflammatory markers in epithelial CF models, suggesting that restoring Ca²⁺ homeostasis may prevent both inflammation and tissue injury in CF.

In this study, using Cftr-deficient zebrafish and human CF bronchial epithelial cells, we *i)* investigated how CFTR dysfunction alters epithelial Ca²⁺ activity to promote inflammation and *ii)* assess the therapeutic potential of Ca^2+^ channel blockers as a news anti-inflammatory strategy in CF. Our work provides *in vivo* evidence that CFTR dysfunction induces abnormal epithelial Ca²⁺ signaling, leading to neutrophilic inflammation in CF. In particular, we identify a pivotal role for L-type VGCC-mediated Ca^2+^ fluxes in driving pathological epithelial Ca^2+^ fluxes, inflammation and tissue damage in CF context. Pharmacological inhibition of L-type VGCCs with the FDA-approved drug verapamil, restored epithelial Ca^2+^ homeostasis, reduced ROS and NF-κB activity, and attenuated neutrophilic inflammation and tissue damage in CF models. These data establish a mechanistic link between CFTR dysfunction, L-type VGCC activity, and epithelial-driven inflammation.

Although evidence suggests that elevated Ca^2+^ levels in CF airway are correlated with excessive production of ROS and pro-inflammatory cytokines, the mechanisms by which CFTR dysfunction alters Ca^2+^ signaling to promote inflammation in CF remain incompletely understood. By generating a transgenic zebrafish line allowing Ca^2+^ flux mapping in epithelia, our findings revealed that CF larvae exhibit abnormal Ca²⁺ elevations in response to injury compared to wild-type controls. Importantly, pharmacological Ca^2+^ signaling-targeting compounds can reduce exuberant neutrophil influx CF animals, confirming that increased epithelial Ca^2+^ signaling contributes to the neutrophilic inflammatory status in CF. In conjunction with data from other CF *in vitro* and *ex vivo* models, these findings provide evidence that CFTR plays an important role in efficiently regulating epithelial Ca^2+^ flux, and establish a link between pathological epithelial Ca²⁺ activity and neutrophilic inflammation in CF.

Importantly, while various Ca²⁺ channels are known to regulate Ca²⁺ responses, and some have been implicated in CF (19,20), our work is the first to implicate L-type VGCCs in CF-related pathological epithelial Ca²⁺ activity. Although classically associated with excitable tissues, L-type VGCCs are also expressed in epithelial and immune cells, where they mediate Ca²⁺ influx during mechanical or inflammatory stimulation. We show that CF-related epithelial Ca²⁺ hyperactivity is also L-type VGCC (Cav1.2)-dependent, as it is abolished by the L-type VGCCs blocker nifedipine. The absence of Cav1.2 transcriptional changes in CF suggests that CFTR influences L-type VGCC activity through post-translational or regulatory mechanisms rather than altered gene expression. Two hypotheses may explain this relationship: *i)* impaired CFTR-mediated Cl⁻ efflux causes membrane depolarization (22), facilitating greater Ca^2+^ entry through activation of VGCCs (23) and *ii)* CFTR dysfunction has been recently associated with elevated activity of protein kinase C (PKC) (24), which may phosphorylate and potentiate L-type Ca^2+^ channel (25). Future studies are needed to dissect these hypotheses.

Although crucial for the regulation of epithelial and immune cell responses, abnormal Ca^2+^ dynamics can drive inflammatory pathologies. Our data support a model in which CFTR dysfunction drives sustained L-type VGCC Ca²⁺ influx, leading to excess ROS generation, NF-κB hyperactivation, uncontrolled neutrophil recruitment, and tissue damage. Importantly, Ca²⁺ dysregulation in CF is not limited to the lung but may contribute to other comorbidities associated with the disease, including CF-related diabetes, gastrointestinal cancers, liver disease and pancreatic fibrosis (2). Sustained Ca²⁺ elevation disrupts mitochondrial function, impairs insulin secretion, induces cellular stress, and accelerates oncogenic signaling, all features observed in CF. These findings position Ca^2+^ signaling as a central driver of both airway inflammation and systemic disease progression, thereby opening broad perspectives for therapeutic intervention.

Overall, these findings identify L-type-mediated epithelial Ca²⁺ signaling as a central driver of CF-related inflammation and tissue damage, highlighting its potential as a therapeutic target. L-type Ca^2+^ channel antagonists are a well-established class of drugs in clinical use for cardiovascular diseases. Among the L-type Ca^2+^ channel antagonists tested (i.e. amlodipine, diltiazem, verapamil), verapamil proved particularly effective, combining anti-inflammatory, pro-resolution, and pro-repair activities. We showed that verapamil efficiently normalized epithelial Ca²⁺ and oxidative activities in both Cftr-deficient zebrafish and human CF epithelial cell line. Thus, by restoring Ca²⁺ / ROS homeostasis, verapamil reduces excessive activation of NF-kB, thereby attenuating neutrophil influx in CF animals.

Unlike diltiazem and amlodipine, verapamil was able to accelerate the resolution of neutrophilic inflammation and promote tissue repair. In CF, chronic neutrophilic inflammation is determined by persistent accumulation of activated neutrophils, along with impaired apoptosis (4, 5). Ca^2+^ signaling is associated with key aspects of neutrophil function and lifespan, including the initiation of apoptosis (17). Although elevated intracellular Ca²⁺ levels have been reported in CF neutrophils, interestingly, in contrast to lymphoid cells, increased cytosolic Ca^2+^ in neutrophils exerts an inhibitory effect on apoptosis (27, 28). While this mechanism remains incompletely defined in CF, restoration of Ca^2+^ homeostasis by verapamil may help re-engage the apoptotic program of CF neutrophils, thereby facilitating a more effective resolution of inflammation.

CF zebrafish exhibit impaired tissue repair after injury, partly due to unresolved inflammation (4, 19). Notably, this defect appears to be Ca²⁺-dependent, as verapamil significantly improved tissue repair outcomes in CF fish. Possible explanations for the pro-repair properties of verapamil in this context include : *i)* its ability to restore epithelial Ca^2+^ homeostasis, a central regulator of regenerative processes including wound closure, cell migration, and extracellular matrix remodeling (29); *ii)* its antioxidant activity may enhances nitric oxide production and upregulates growth factors such as VEGF, both impaired in CF (30), thereby promoting angiogenesis and tissue repair (31); *iii)* by blocking L-type VGCCs, verapamil prevents oxidative damage, dampens NF-κB activation, and limits neutrophilic inflammation, collectively creating a permissive environment for tissue healing and repair. Consistent with this, verapamil has been shown to alleviate oxidative stress and suppress NF-κB-mediated inflammatory gene expression in a mouse model of inflammatory arthritis, ultimately reducing tissue injury (32).

Ideally, host-modulatory therapies should promote the resolution of inflammation while minimizing the risk of impeding antimicrobial defenses. The anti-inflammatory proprieties of verapamil raise critical questions regarding its safety in the context of chronic bacterial infections, which are highly prevalent in CF. Importantly, our data show that verapamil selectively attenuates pathological inflammation without impairing neutrophil chemotaxis per se towards fMLP, in contrast to other host-modulatory compounds we have recently evaluated, such as roscovitine, which, by reducing epithelial oxidative responses, compromises antimicrobial defense and, consequently, exacerbates infections (33). This selectivity suggests that verapamil can rebalance host responses while preserving essential immunity. Nonetheless, studies in *in vivo* infection models are required to confirm its safety in the context of chronic airway infections. Beyond its immunomodulatory effects, verapamil has been shown to increase antibiotic efficacy; by inhibiting bacterial efflux pumps, potentiates the antimicrobial activity of bedaquiline (34) and clofazimine (35), two drugs with efficacy against *M. abscessus* (36), and other drugs targeting *P. aeruginosa* and *S. aureus* (37, 38). Such dual benefits, inflammation control and improved antimicrobial efficacy, would be particularly valuable in CF, where multidrug-resistant infections are common.

Given its long-standing clinical use, favorable safety profile, oral bioavailability, well-characterized pharmacokinetics, and low cost, verapamil represents a strong candidate for repurposing in CF. These features could facilitate the design of pilot clinical trials to assess verapamil in CF patients, focusing on safety, tolerability, airway inflammation, lung function, and exacerbation frequency. Importantly, because persistent inflammation occurs even in patients receiving CFTR modulators, verapamil could provide an adjunct option, expanding treatment strategies for individuals not eligible for, or not fully benefiting from, CFTR modulators.

To conclude, our findings identify L-type VGCCs as central mediators of Ca²⁺-driven inflammatory damage in CF and support repurposing of VGCC antagonists, particularly verapamil, as candidate anti-inflammatory therapies in CF.

## Acknowledgment

We thank Etienne Meunier for his discussions and advice, as well as all the people that provide valuable reagents for this study. We acknowledge the ZEFIX-LPHI Aquatic model facility (ZEFIX, Biocampus, University of Montpellier), and Catherine Gonzalez and Victor Goulian for zebrafish maintenance and care. We thank the Montpellier GenomiX quantitative qPCR facility of the University of Montpellier, and Philippe Clair for his valuable assistance. Finally, we thank the University of Montpellier, the University of Toulouse and the University of Sheffield for support.

