## Supplementary figures and images for "Verapamil limits inflammation by restoring VGCC-driven epithelial Ca^2+^ in models of cystic fibrosis"

### supp figures

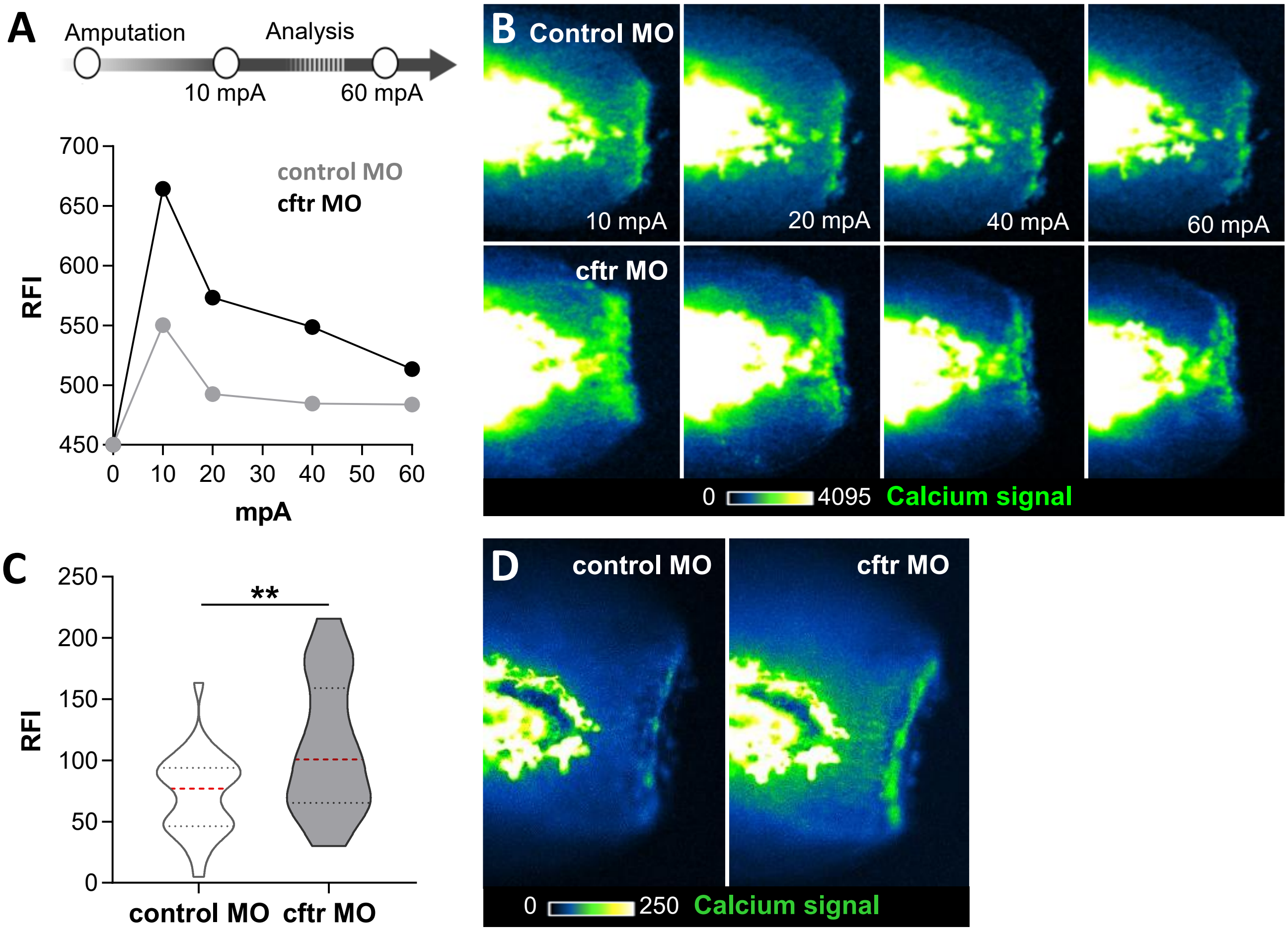

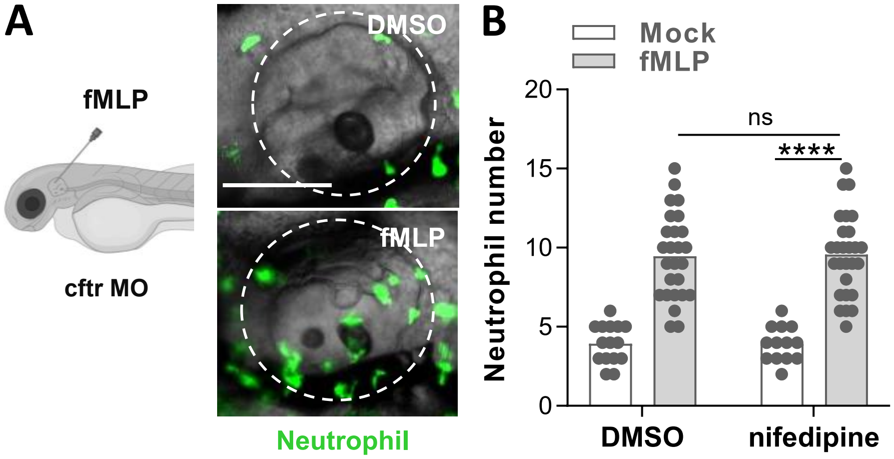

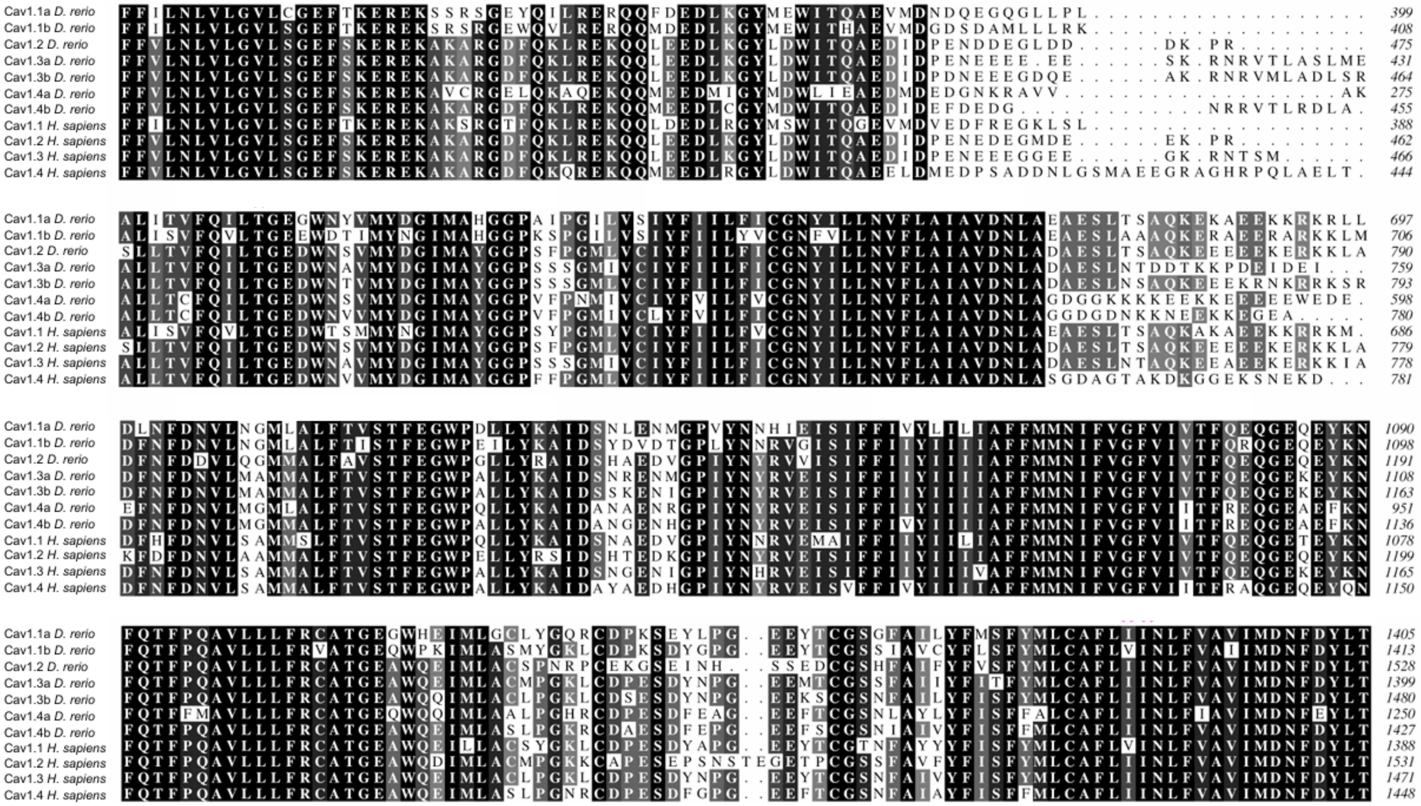

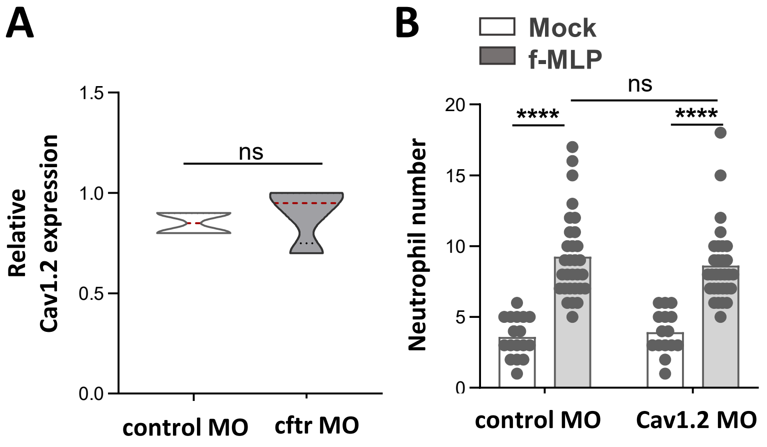

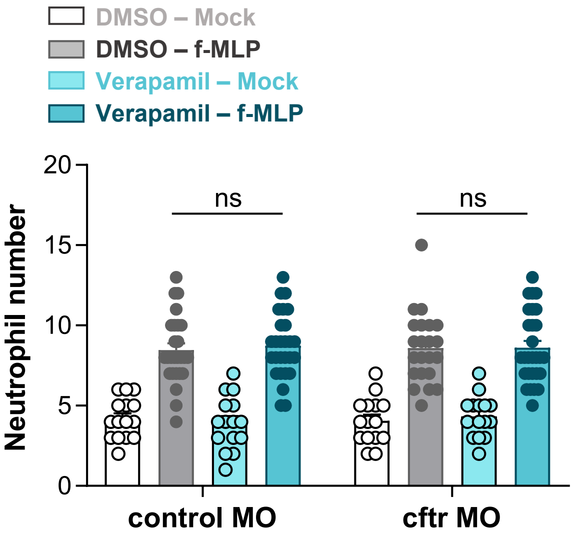

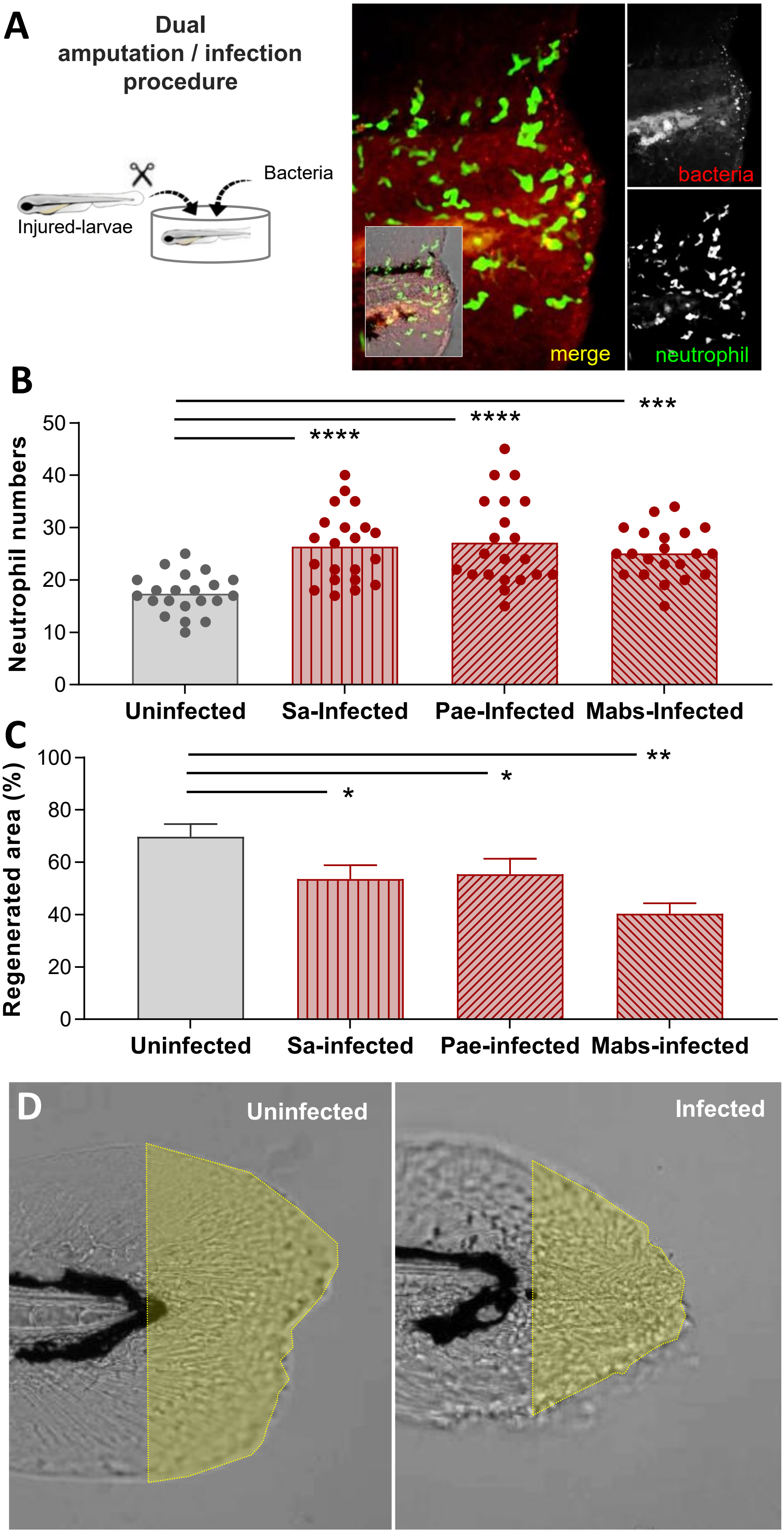

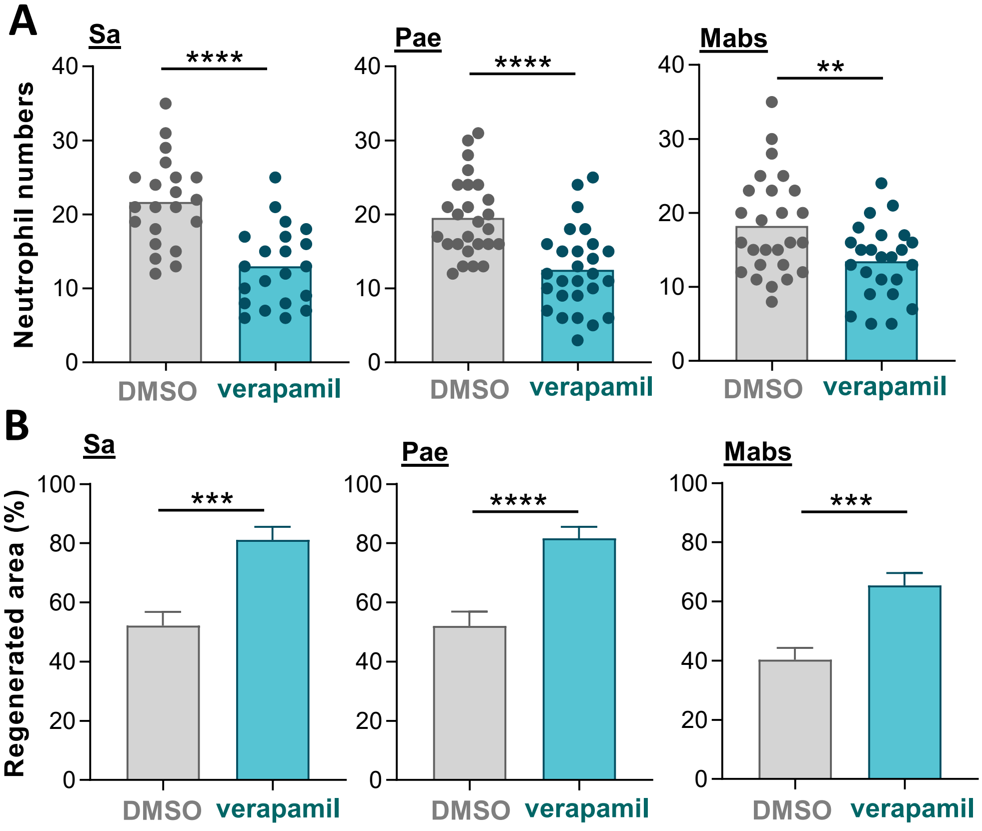

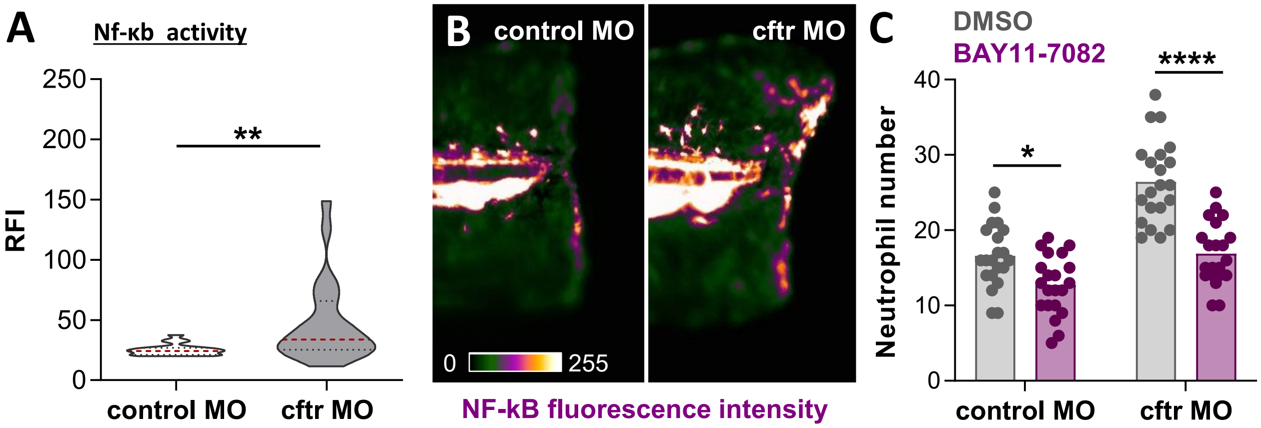
