## Supplementary material for "Verapamil limits inflammation by restoring VGCC-driven epithelial Ca^2+^ in models of cystic fibrosis": Supp data

Sarahdja Cornélie^1^, Bastien Suire^2^, Sylvaine Huc-Brandt^1^, Laure Yatime^1^, Christian M Dupont^1^, R Andres Floto^3,4^, Stephen A Renshaw^5^, Georges Lutfalla^1^, Céline Cougoule^2^, Audrey Bernut^1#^

^1^ Laboratory of Pathogens and Host Immunity, CNRS, Inserm, University of Montpellier, Montpellier, France.

^2^ Institute of Pharmacology and Structural Biology, CNRS, University of Toulouse, Toulouse, France.

^3^ Molecular Immunity Unit, University of Cambridge Department of Medicine, MRC-Laboratory of Molecular Biology, Cambridge, United Kingdom.

^4^ Cambridge Centre for Lung Infection, Royal Papworth Hospital, Cambridge, United Kingdom.

^5^ Bateson Centre, School of Medicine and Population Health, University of Sheffield, Sheffield, United Kingdom.

^#^ correspondence:/ Laboratory of Pathogen and Host Immunity, CNRS, Inserm, University of Montpellier, 34090 Montpellier, France.

**SUPPLEMENTARY METHODS**

**Zebrafish Husbandry and Ethics Statement**

Zebrafish (*Danio rerio*) work was conducted by following the 3Rs -Replacement, Reduction and Refinement- principles in compliance with the European Union guidelines for handling of laboratory animals. Adult zebrafish were maintained in the ZEFIX-Lphi aquaria (CNRS, Inserm, University of Montpellier, Montpellier, France; license number C34-172-37), according to the local animal welfare standards protocols approved by the Direction Sanitaires et Vétérinaires de l’Hérault, the Comité d’Ethique pour l’Expérimentation Animale de la région Languedoc Roussillon and the French Ministry of Agriculture and Food.

Zebrafish experiments were carried out using the wild type AB line or the golden mutant. To study neutrophil response during inflammation, the *TgBAC(mpx:eGFP)^i114^* reporter line was used (1). The *Tg(il1b:eGFPF)^ump3tg^* and *Tg(pNF-κB:EGFP)^sh235^* reporter lines were used to visualize the transcriptomic expression of *il1b* and *nf-κb* reciprocally (2,3).

All experimental procedures were performed by authorized staff on larvae prio to the fish-free feeding stage (< 5 days post-fertilization (dpf)). Zebrafish eggs were obtained from pairs of adult fish by natural spawning and raised in E3 water (5 mM NaCl, 0.17 mM KCl, 0.33 mM CaCl2, 0.33 mM MgSO4) at 28°C and exposed on a 14:10 hours light/dark cycle to maintain proper circadian conditions. The number of animals used for each procedure was guided by pilot experiments or by past results. For zebrafish anesthesia procedures, larvae are immersed in a 168 mg/L tricaine (Sigma-Aldrich) or 0.0075 % eugenol (Fisher Scientific) solution in E3 water. When required, larvae were euthanized using an overdose of tricaine (500 mg/L).

**Generation of an epithelial calcium reporter zebrafish**

The promoter of type I keratin, krtt1c19e, a marker of the basal epidermal layer used to drive the specific expression of the GCaMP6 Ca^2+^ indicator in epithelial cells, was amplified from pTol2Mfap4:GCaMP6 (4) using primers (5’-TCCCTCGTTGTCAGTCTTCT-3’) and (5’-TATAGCGGCCGCCCATGTGGATGGTGGTTGGT-3’). The 3.3 kb amplified fragment was digested by NotI, and ligated to the coding phase of the GCaMP6 protein on a I-SceI meganuclease and Tol2-derived vector (GenBank accession no. GU394080). The resulting plasmid was injected, together with the I-SCe1 meganuclease, into one-cell stage embryos. F0 injected embryos were screened by exposure to 5 µM ionomycin in Ringer’s solution, which stimulates Ca^2+^ entry into cells, and a stable line was established.

**Morpholino Injections**

Morpholinos used in this study were purchased from Gene Tools. The splice-blocking morpholino for *cftr* knock-down (5’-GACACATTTTGGACACTCACACCAA-3’) was prepared and injected into one-cell-stage zebrafish as previously described (5). Two splice-blocking morpholinos for cacna1c knock-down (5’-CCCGTTCCTAGACAGACGAAACAGA-3’) and (5’-GGATCTTGCACTCACCTACGACCA-3’) were prepared and injected into one-cell-stage zebrafish as described (6). A standard control morpholino (5'-CCTCTTACCTCAGTTACAATTTATA-3') was used as a negative control.

**Drug Treatments of Zebrafish larvae**

To explore the therapeutic efficacy of FDA-approved L-type VGCC antagonists, diltiazem (10 µg/ml), amlodipine (10 µg/ml) or verapamil (20 or 40 µg/ml) were added in zebrafish water. Nifedipine (10 µM) was used to inhibit L-VGCC. Capsazepin was used to inhibit TRPV (5 µM)*,* Cpd5J-4 was used to inhibit SOCE (5 µM). For all Ca^2+^ modulators, the doses that we identified did not generate toxicity. Dimethyl sulfoxide (DMSO) was used as vehicle control.

**RNA Isolation from Zebrafish larvae and qRT-PCR Analysis**

Total RNA from a pool of 10-15 larvae per biological experiment was extracted using Nucleospin RNAII kit (Macherey-Nagel) at time points indicated in Figure legends and cDNA synthesized with M-MLV reverse transcriptase (Invitrogen). Real-time RT-PCR were performed using SensiFAST SYBR Green No ROX mix (Thermo Fisher Scientific) on a LightCycler 480 instrument (Roche) as described (7) and gene expressions were detected with the gene-specific primers provided in Table 2. Each experiment was run in triplicate. ΔCT was calculated using the housekeeping gene ef1α as a reference gene. Relative expression levels were calculated using the 2−ΔCt method.

***In vivo* Inflammation Assays**

Inflammation was induced by distal or proximal tail fin amputation on 3 days post-fertilization (dpf) anesthetized larvae using a microscalpel (5 mm depth; World Precision Instruments) according to established procedures (8). Neutrophil response was observed and evaluated by manually assessing the number of cells at wound sites at relevant time points using an Olympus MVX10 fluorescence microscope (Olympus, Life Science) equipped with a X-Cite Xylis LED (Excelitas Technologies) light source as previously defined (8): 3 or 4 hours post-amputation (hpA) (acute phase of neutrophil response) or 8 hpA (resolution phase of neutrophil response).

Epithelial Il1b and Nf-κb signaling at the wound sites (50 µm anterior from the wound marge) were observed and captured at 2, 3 or 4 hpA, using a fluorescence microscope (Olympus, Life Science). Images were acquired with a Hamamatsu ORCA-spark Digital CMOS C11440-36U camera (Hamamatsu Photonics Inc) and processed using Olympus CellSens Standard 3.1 software (Olympus, Life Science). Epithelial Il1b and Nf-κb signaling were assessed in Fiji (ImageJ software) using the intensity of fluorescence and normalized to uninjured animals (control MO or *cftr* MO; untreated or treated larvae).

**Neutrophils Recruitment Assays**

The f-Met-Leu-Phe (fMLP, Sigma-Aldrich) was used to induce neutrophil chemotaxis pharmacologically, as previously described (8). Briefly, fMLP was injected into the otic vesicle of 3 dpf larvae. Neutrophil response was observed and evaluated by manually assessing the number of cells at the injection sites using a fluorescence microscope (Olympus, Life Science).

***In vivo* Calcium Activity Assay**

Epithelial Ca^2+^ response post-injury was detected using Fluo-3 AM green (Thermo Fisher Scientific) and the *Tg(krtt1c19e:GCaMP6)^ump14Tg^* reporter line generated in this study.

*Fluo-3 AM method*/ Anesthetized zebrafish larvae were tail fin injured and immediately soaked in Fluo-3 AM reagent (10 µM) during 30 min. Injured fish are rinsed twice and incubated in fish water for 30 minutes. Epithelial Ca^2+^ response was observed and measured at 1 hpA using a fluorescence microscope (Olympus, Life Science) and then processed and analyzed with Fiji (ImageJ software) using the intensity of fluorescence.

*Transgenic line method/* Anesthetized larvae were amputated were tail fin injured, then epithelial Ca^2+^ response was observed and measured at 10, 20, 40 or 60 mpA using a fluorescence microscope (Olympus, Life Science) and then processed and analyzed with Fiji (ImageJ software) using the intensity of fluorescence.

***In vivo* Oxidative Activity Assay**

Epithelial oxidative response post-injury was detected using CellROX® green (Thermo Fisher Scientific) following established protocols (9). Briefly, living zebrafish larvae were soaked in CellROX® reagent prior tail fin amputation procedure during 30 minutes, then ROS production at the wound sites (50 µm anterior from the wound marge) were observed and captured at 30 mpA using a fluorescence microscope (Olympus, Life Science). Epithelial oxidative response was assessed in Fiji (ImageJ software) using the intensity of fluorescence and normalized to uninjured animals (control MO or *cftr* MO; untreated or treated larvae).

***In vivo* Tissue Repair Assay**

Tissue repair assay was performed following established methods (10). Briefly, 2 dpf embryos were anesthetized then tail fins amputated at the boundary of the notochord without injury to the notochord (distal injury). Tissue repair performances were evaluated by assessing the regenerated tail fin area at 3 dpA under a microscope (Olympus, Life Science). Images were processed and analyzed with Fiji (ImageJ software): regrowth area was measured by outlining the total fin tissue area distal to the notochord using the polygon tool. Percentage of regeneration was calculated by normalizing the regenerated tail fin areas versus fin areas of unamputated animals (control MO or *cftr* MO; untreated or treated larvae).

**Bacterial strains, growth conditions and tail wound infection challenges**

*Staphylococcus aureus* (SH1000 strain carrying pMV158-mCherry (11), generously provided by Simon Foster (Florey Institute, Sheffield, UK)), *Pseudomonas aeruginosa* (PAO1 strain) and *Mycobacterium abscessus* sensu stricto (CIP104536T strain, morphotype smooth (S)), were used for this study. *S. aureus* were grown using Brain Heart Infusion (BHI) media (MilliporeSigma) with tetracycline 5 µg/ml (Sigma-Aldrich). *P. aeruginosa* were grown using Luria-Bertani media (LB Miller, MilliporeSigma). *M. abscessus* were grown in Middlebrook 7H10 Agar and Middlebrook 7H9 Broth media (MilliporeSigma), supplemented with 10% acid/albumin/dextrose/catalase (ADC) enrichment (MilliporeSigma). To prepare *S. aureus* and *P. aeruginosa* inoculates, 1 ml of appropriate medium was inoculated with a fresh colony of bacteria and incubated at 37°C overnight with shaking. 500 µl of this overnight culture was then added to 10 ml of appropriate medium and incubated at 37°C with shaking to achieve growth to mid-logarithmic phase (OD600 ≈ 0,6-0,8). *M. abscessus* inoculates were prepared as previously described (12). Next, to generate infected medium, bacteria were harvested by centrifugation, washed with Phosphate-Buffered Saline (PSB, GibcoTM, Thermo Fisher Scientific), and resuspended at an OD600 of 1 in sterile water. Caudal tail fin amputation of anesthetized-larvae was performed as described above, then injured larvae were immediately transferred in sterile or infected medium for 1 hour at 28°C. Uninfected and infected larvae were then rinsed three times with sterile wtaer to wash away bacteria, and maintained at 28°C until imaging for neutrophil mobilization or wound healing assays as described above.

**Minimum Inhibitory Concentration**

Determination Minimum Inhibitory Concentrations (MICs) were determined using the microdilution method, in cation-adjusted Mueller-Hinton broth, according to the Clinical and Laboratory Standards Institute (CLSI) guidelines (13). Serial 10-fold dilutions of log-phase cultures were plated and incubated at 37°C for 3 to 4 days, and the MICs were recorded by visual inspection and defined as the minimum concentration required to inhibit 99% of the growth. Experiments were done in triplicate in three independent occasions.

***In vitro* inflammation assays and treatments in human bronchial epithelial cells**

To measure calcium activity, human bronchial epithelial cells (CFBE41o- and 16HBE14o-) were incubated at 37°C for 30 min with 5 µM of Fluo 3-AM, washed twice with Ca^2+^-free Hank's Balanced Salt Solution (HBSS), then further incubated for 15 min at 37°C in the same buffer to allow complete de-esterification of the dye. Next, cells were pre-treated with 25 µM verapamil for 30 min at 37°C, stimulated with 1 µM ionomycin, and then Fluorescence kinetics were recorded every 30 s for 2 h (240 cycles). Calcium activity was quantified as relative luminescent units (RLU).

To measure oxidative activity, cells were incubated at 37°C for 30 min with CellROX. Cells were pre-treated with 25 µM verapamil for 30 min at 37°C, stimulated with 25 µM cumene hydroperoxide, and then fluorescence kinetics were recorded every 30 s for 2 h. Oxidative activity was quantified as RLU.

**Quantification and Statistical Analysis**

Statistical analyses were performed using Prism 10.0 (GraphPad Software) and are detailed in each figure legend: ns, not significant (*P*≥0.05); **P*<0.05; ***P*<0.01; ****P*<0.001; and *****P*<0.0001.

**SUPPLEMENTARY DATA**

**Figure S1. Cftr-depleted zebrafish exhibit increased epithelial Ca^2+^ fluxes to injury. (A-B)** Control (control MO) and *cftr* morphants (cftr MO) *Tg(krtt1c19e:GCaMP6))^ump14Tg^* were tail fin amputated, then epithelial Ca^2+^ activity at wound sites was measured and analyzed every 10 minutes during 1 hour. Changes in epithelial Ca^+2^ fluxes at wound sites in single individuals, showing representative relative fluorescence intensity (RFI) in the control vs CF animals **(A)**, and associated pseudocolored photomicrographs of injured tails revealing epithelial Ca^2+^ activity **(B)**. **(C-D)** Control MO and cftr MO were tail fin amputated and immediately stained with Fluo-3 AM to label and analyze Ca^+2^ signaling. Relative epithelial Ca^+2^ activity at the wound sites at 1 hpA (n=30 from 3 independent experiments; Student T-test) **(C)**, and associated pseudocolored photomicrographs of injured tails **(D)**.

**Figure S2. Nifedipine exposure does not affect neutrophil chemotaxis *per se* in zebrafish. (A)** fMLP or DMSO (as control) was injected into the otic cavity of cftr MO *TgBAC(mpx:eGFP)^i114^*. Representative photomicrographs of fMLP-injected animal compared to DMSO-injected control. fMLP induces neutrophil mobilization to the simulated site. **(B)** Neutrophil mobilization into the otic cavity in response to DMSO or fMLP injection in *TgBAC(mpx:EGFP)i114* larvae treated or not with nifedipine (3hpA, n=15-30 from three independent experiments; Two-way ANOVA, Tukey’s multiple comparisons test).

**Figure S3. Analysis of the sequence conservation between human Cavs and zebrafish Cavs.** Protein sequences of human Cavs and zebrafish Cavs were aligned using Clustal Omega from the EMBL-EBI server (14). Analysis of the sequence conservation was done with ALINE (15).

**Figure S4. Genetic inhibition of Cav1.2 does not affect neutrophil chemotaxis *per se* in zebrafish. (A)** mRNA levels of *Cav1.2* at 2 hpA determined by RT qPCR (relative gene expression from four independent experiments performed in triplicates, Mann-Whitney U test). **(B)** Control MO and Cav1.2 MO *TgBAC(mpx:eGFP)i114* were injected with f-MLP or DMSO into the otic vesicle, then the number of neutrophils mobilized at injection site was observed and counted at 3hpA (n=15-30 from three independent experiments; Two-way ANOVA, Tukey’s multiple comparisons test).

**Figure S5. Verapamil treatment does not affect neutrophil chemotaxis *per se* in zebrafish.** Neutrophil mobilization into the otic cavity in response to DMSO or fMLP injection in control MO and *cftr* MO *TgBAC(mpx:EGFP)i114* larvae treated or not with verapamil or DMSO (3hpA, n=15-27 from 3 independent experiments; Two-way ANOVA, Tukey’s multiple comparisons test).

**Figure S6. Infections worsen inflammation and repair defects in CF zebrafish. (A-B)** *cftr* MO *TgBAC(mpx:EGFP)^i114^* were injured and infected with fluorescent *S. aureus* (Sa), *P. aeruginosa* (Pae) or *M. abscessus* (Mabs). (B) Neutrophil number at the infected wound was counted at 2 hpA (3 independent experiments, One-way ANOVA with Dunnet's multiple comparisons test). **(C-D)** *cftr* MO were injured and infected with *S. aureus, P. aeruginosa* or *M. abscessus* for 4 hours. The potential of tissue repair was evaluated by measuring regenerated fin areas at 3 dpA (n=21 from 3 independent experiments, One way ANOVA with Dunnet's multiple comparisons test).

**Figure S7. Verapamil reduces neutrophilic inflammation and tissue injury in CF zebrafish during bacterial infections. (A)** *cftr* MO *TgBAC(mpx:EGFP)^i114^* were pre-treated with verapamil or DMSO prior to caudal fin amputation procedure, then injured, and immediately exposed with *S. aureus, P. aeruginosa or M. abscessus* for 1 hour. Double injured/infected larvae were then put back in treatments until analysis. **(A)** Neutrophil number at the infected wound counted at 2 hpA (3 independent experiments, Two-tailed Student t-test). **(B)** *cftr* MO were pre-treated with verapamil or DMSO prior to caudal fin amputation procedure, then injured, and immediately exposed *with S. aureus, P. aeruginosa* or M. abscessus for 1 hour. Double injured/infected larvae were then put back in treatments for 4 hours. Regenerated fin areas were evaluated 3 dpA (n=14-18 from 2 independent experiments, Mann-Whitney U test or Two-tailed Student t-test).

**Figure S8. Exuberant wound-induced NF-κB signaling drives the overactive neutrophil response in CF zebrafish. (A-B)** NF-κB activity upon wounding in control and cftr MO *Tg(pNF-kB:EGFP)^sh235^*. Control and cftr MO *Tg(pNF-kB:EGFP)^sh235^* were caudal fin amputated then the activity of NF-κB was observed and analyzed at 2 hpA. Relative NF-κB intensity as violin plots (n=20-22 from 3 independent experiments; Mann-Whitney U test) **(A)** and associated pseudocolored photomicrographs of injured tails **(B)** revealing NF-κB activity at the wound-edge. Loss of CFTR leads to an increase in NF-κB activation in response to injury. **(C)** Inhibition of NF-κB was carried out using BAY11-7082. Neutrophil number at the wound at 4 hpA (3 independent experiments, Two-way ANOVA with Tukey’s multiple comparisons test).
